# Possible Anti-viral effects of Neem (*Azadirachta indica*) on Dengue virus

**DOI:** 10.1101/2020.04.29.069567

**Authors:** Vikas B Rao, Kalidas Yeturu

## Abstract

Dengue virus (DENV) has become a major health threat worldwide accounting for 50-100 million infections every year and keeping 2.5 billion people at a risk of the infection. Seriousness of the viral infection can be attributed to its lethality when not treated in time and potential to cause health complications post infection. Currently there are only preventive strategies and development of vaccination is still in its infancy of research. It is therefore highly necessary to discover newer drugs and therapies for this deadly virus. In this paper we report important insights we have obtained through a computational analysis of small molecules of Neem (Azadirachta indica) against dengue viral proteins and its required proteins in human. Our study involves identification of the effect of specific small molecules of Neem on proteins of human and virus corresponding to different pathways using simulated molecular binding analyses. We report here Gedunin and Pongamol contained in naturally occurring Neem as potential drugs against the Dengue virus.

**Significance Statement:** We report important ligands in Neem that have potential antiviral activity against Dengue. Our selection of Neem for testing for antiviral properties has been inspired from Ayurveda. Due to unhygienic living conditions that facilitate mosquito breeding, Dengue is a major threat in developing countries causing millions of deaths. Despite the severity of the infection, no specific antiviral drug is available. The results obtained, in terms of newer potential ligands against Dengue are significant as this provides a basis for experimentally verifying and extending the same to develop a cure.We hope that this study would would spur scientific curiosity and undertaking of further elaborate experimental studies.

---

**D**engue virus (DENV) is a single positive stranded, arthropod-borne virus, of the Flaviviridae family. All of the four identified serotypes (DENV 1 - 4) are highly infectious in the human host [11]. According to a WHO report (as of 2013-July-14), a total of 2.5 billion people are at a risk of infection with over 50-100 million infections reported every year worldwide (http://www.who.int/mediacentre/factsheets/fs117/en/). *Aedes aegypti* and *Aedes albopictus* are the mosquito vectors, through which DENV spreads [4]. The incubation period of the virus in the host varies from individual to individual ranging from 4 to 7 days post infection [12]. If the infection is not diagnosed and treated during its early stages, it causes Dengue haemorrhagic fever (DHF) and Dengue shock syndrome (DSS) which are often fatal [9]. DENV has a small genome encoding for ten proteins [7] and it interacts with a number of human proteins [7, 1]; the details of which are provided in the Materials and Methods section. Currently, there are only preventive strategies to reduce the incidence of infection by disinfecting the mosquito breeding grounds. Research on development of vaccine is still in its infancy. It is therefore necessary to explore newer drugs which can reduce the viral loads during infection. Prior *in vitro* research has shown that Neem inhibits RNA synthesis in DENV thus arresting replication of the virus and hence the infection [10]. Given that a biological system can be studied at various levels of abstraction, we have carried out molecular level analysis here to determine ligands of Neem that may block human proteins required by the virus and viral proteins themselves. We report here some of the Neem ligands (Gedunin and Pongamol primarily) as possible drugs against the virus based on molecular binding simulation *in silico* using Autodock [8].

## Materials and Methods

The approach we have followed here is to determine the effect of small molecules present in Neem on the proteins of human and DENV through the use of molecular binding simulation software, Autodock [8]. We have taken here structures of small molecules on one hand and structures of the proteins of human and virus on the other hand and carried out a molecular docking exercise. The structures of proteins are obtained from the Protein Data Bank [2]. The structures of ligands associated with Neem such as Azadirachtin, Nimbin, Nimbidin, Salanin, Pongamol, Meliantriol, Meldenin, Gedunin, and a list of Tannins (Catechin, Epicatechin, Epigallocatechin, Gallic acid and Gallocatechin) [6, 3] were obtained from the Zinc database [5]. There were reports of Gedunin, Nimbolide, Azadirachtin and Mahmoodin exhibiting anti-malarial properties as well [3, 5]. Our efforts here are to examine effects of many a ligand of Neem at the molecular structural level on the proteins of virus and those virus-required proteins of human. The proteins encoded by DENV genome are namely capsid protein (C), envelope protein (E), and seven non-structural proteins NS1, NS2A, NS2B, NS3, NS4A, NS4B and NS5 [7]. Important human proteins with which virus interacts are the immune system molecules such as Interleukin (IL-5) and Signal transducer and activator of transcription 2 (STAT2) among others. The details of the protein names along with their corresponding PDB structure codes are indicated in Table 1, (1a for human and 1b for DENV proteins). A list of important human and dengue virus proteins that interact [7] is provided in table 2. For each of the proteins, its PDB structure is considered along with each of the ligand structures and a molecular binding simulation is carried out using Autodock [8]. Autodock software is free for academic research and is highly popular. It employs a genetic algorithm to encode torsional space of a ligand and generates many possible conformations of the small molecule binding to various sites on the structure of a given protein molecule. Infeasible energy conformations such as bumping of atoms of proteins and ligand are removed and only feasible conformers of the ligand are output along with their binding energy values. The parameter settings and output results on docking run of each of the Neem ligands (8 ligands) against each of the PDB structures of proteins (21 structures) are provided in supplementary material (S1). For each docking exercise, protein centre and grid volume are chosen interactively using Autodock-tools and default parameters are set.

**Table 1:** Human proteins and their PDB accession code

| Protein | PDB Code | Protein | PDB Code |
| --- | --- | --- | --- |
| **1(a) - Human Proteins** |  |  |  |
| CD14 | 4GLP | Endoplasmin GRP94 | 1U20 |
| CLEC5A | 2YHF | FC gamma receptor | 1H9V |
| Coronin | 2AKF | Beta Haemoglobin | 3ONZ |
| DAXX sub7 | 4H9S | Heat Shock Protein 90 | 1QZ2 |
| DC-SIGN | 1XAR | IL-5 | 3VA2 |
| Laminin receptor | 3BCH | Myosin | 2LV6 |
| **1(b) - DENV Proteins** |  |  |  |
| Capsid Heterotetramer (C) | 3J2P | (4) NS3 | 2JLQ |
| (1) Envelope 3 (E3) | 3IRC | (1) E3 Post-fusion | 3G7T |
| (1) NS2B/NS3 Protease | 3L6P | (1) NS2B/NS3 Protease mutant | 3LKW |
| NS3 RDRP | 4HHJ | NS5 RDRP | 2J7U |
| NS3 Protease Helicase | 2FOM |  |  |

Legend - Protein name and 4 letter PDB codes are indicated.

**Table 2:** Human protein - DENV protein interaction

| DENV protein(s) | Human protein(s) |
| --- | --- |
| NS5 | STAT2 (2KA4) |
| NS2A, NS4A, NS4B | IL-5 (3VA2) |
| NS2B/NS3 Protease | Endoplasmin (1U20) |
| Capsid | DAXX (4H9S, 4H9N, 4H9Q) HBB (3ONZ) |

## Results and Discussion

Autodock [8] was used to carry out docking of the Neem ligands against the proteins of DENV and human which are involved in the processes of infection. Autodock performs molecular binding simulation and outputs a number of ligand conformers binding to cavities on the surface of a protein along with computed energies released upon binding. The highest of energy released upon binding for each of the ligands to the selected proteins is tabulated in tables 3 and 4 for human proteins and viral proteins respectively. Analysis of the binding energies has shown that Gedunin is the most potential binder to a number of human and viral proteins alike. DENV interferes with the functioning of the host by binding to various receptor sites and upregulating or down-regulating the functioning of the host proteins. DENV viral entry can be arrested through blocking of the receptor sites on the surface of human cells, leaving no vacant receptor sites for DENV binding to happen. The results corroborate the same theory. We report here that Gedunin binds efficiently to human proteins CD14, DAXX, DC-SIGN and IL-5 which are present on the human cell surface facilitating the attachment of DENV ultimately resulting in viral entry and infection [7, 1].

**Table 3:**
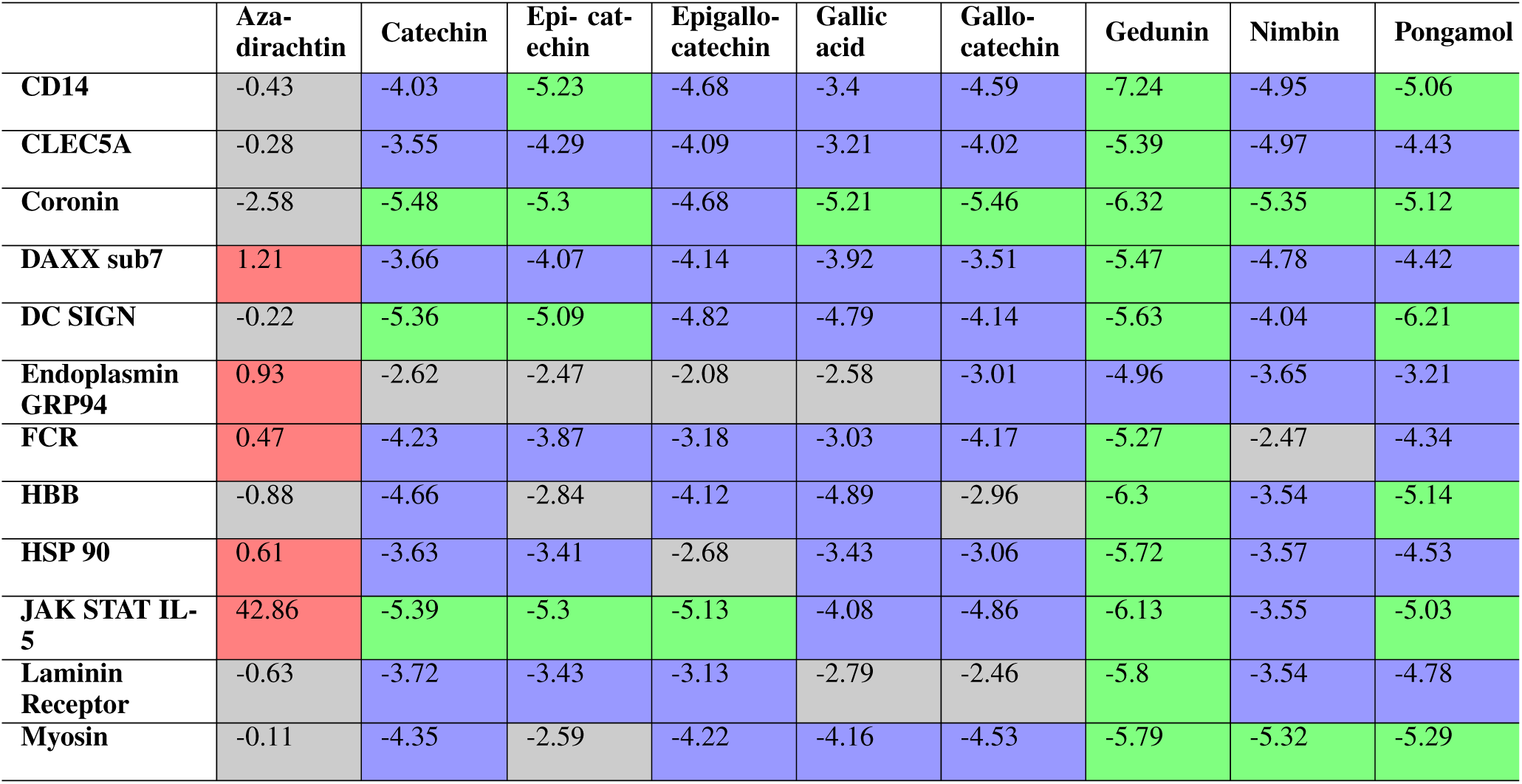
Binding energy values of human proteins against Neem ligands

|  | Aza-dirachtin | Catechin | Epi- catechin | Epigallo-catechin | Gallic acid | Gallo-catechin | Gedunin | Nimbin | Pongamol |
| --- | --- | --- | --- | --- | --- | --- | --- | --- | --- |
| CD14 | -0.43 | -4.03 | -5.23 | -4.68 | -3.4 | -4.59 | -7.24 | -4.95 | -5.06 |
| CLEC5A | -0.28 | -3.55 | -4.29 | -4.09 | -3.21 | -4.02 | -5.39 | -4.97 | -4.43 |
| Coronin | -2.58 | -5.48 | -5.3 | -4.68 | -5.21 | -5.46 | -6.32 | -5.35 | -5.12 |
| DAXX sub7 | 1.21 | -3.66 | -4.07 | -4.14 | -3.92 | -3.51 | -5.47 | -4.78 | -4.42 |
| DC SIGN | -0.22 | -5.36 | -5.09 | -4.82 | -4.79 | -4.14 | -5.63 | -4.04 | -6.21 |
| Endoplasmin GRP94 | 0.93 | -2.62 | -2.47 | -2.08 | -2.58 | -3.01 | -4.96 | -3.65 | -3.21 |
| FCR | 0.47 | -4.23 | -3.87 | -3.18 | -3.03 | -4.17 | -5.27 | -2.47 | -4.34 |
| HBB | -0.88 | -4.66 | -2.84 | -4.12 | -4.89 | -2.96 | -6.3 | -3.54 | -5.14 |
| HSP 90 | 0.61 | -3.63 | -3.41 | -2.68 | -3.43 | -3.06 | -5.72 | -3.57 | -4.53 |
| JAK STAT IL-5 | 42.86 | -5.39 | -5.3 | -5.13 | -4.08 | -4.86 | -6.13 | -3.55 | -5.03 |
| Laminin Receptor | -0.63 | -3.72 | -3.43 | -3.13 | -2.79 | -2.46 | -5.8 | -3.54 | -4.78 |
| Myosin | -0.11 | -4.35 | -2.59 | -4.22 | -4.16 | -4.53 | -5.79 | -5.32 | -5.29 |

**Table 4:**
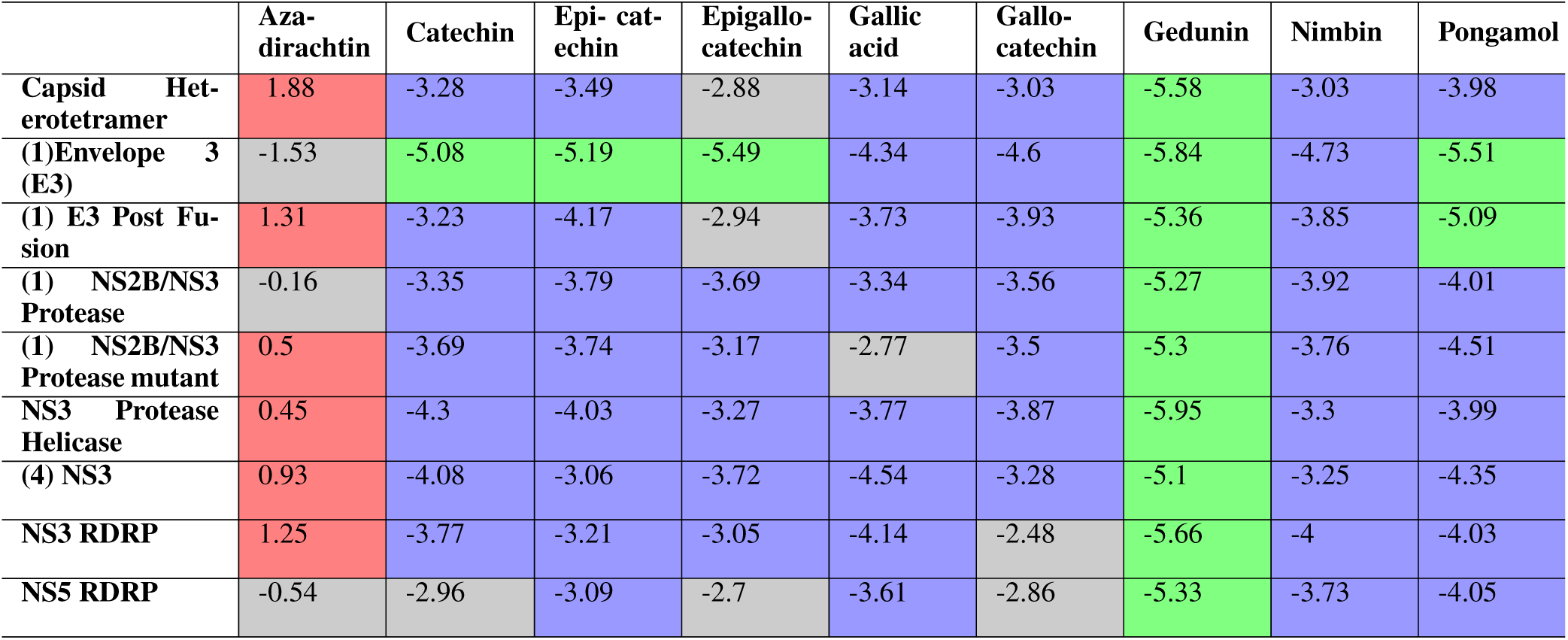
Binding energy values of DENV proteins against Neem ligands

We have specifically analyzed for binding of Gedunin to JAK/STAT Human Interleukin 5 protein of the host which is important in binding of the virus to the host cell surface [7]. The docking simulation reports an energy of 6.13kcal/mol released upon binding of Gedunin to the protein. A careful examination of the binding site of the protein along with the ligand after docking exercise has revealed that such a binding is indeed possible due to chemical complementarity and possibility of a hydrogen bond between oxygen atom of the ligand and a nitrogen atom of the ARG266 residue of the protein at the binding site (Figure 1). The binding also indicates that structurally the ligand is well accommodated in the pocket occupying the space aptly. It is therefore indicative of a possibility of real binding of Gedunin to the protein *in vitro*. In addition, we have observed high binding affinities of Gedunin for DENV proteins namely NS3 RNA polymerase, NS3 protease helicase, capsid and envelope proteins, which are very important for biological functioning of the virus. The capsid and envelope proteins are of high importance as they mediate the entry of the virus into the host cell while the RNA polymerase (NS3 and NS5) operate for synthasis of Dengue proteins inside the host cell [7]. The next better ligand after Gedunin is Pongamol having higher binding affinities compared to the rest of the ligands. Human IL-5 protein that Gedunin and Pongamol bind to correspond to the important JAK/STAT pathway of the immune system which is targeted by DENV proteins namely NS2A, NS4A and NS4B [7]. Gedunin, Pongamol and other ligands as well, are portraying high binding affinities to the human protein DC-SIGN, which is considered the most important DENV receptor mediating the entry of the virus into the host [1]. The above mentioned ligands bind to the common receptor sites preventing DENV-human interaction at the molecular level through blocking of receptor sites in human. The computational study and results we report here would therefore be very important in considering the small molecules of Neem for planning further experimental analyses. One immediate future direction we see here is the characterization of pharmacodynamic behaviour of the small molecules of Neem and systems biological study of involved host proteins. Another direction of research would be to consider a general case of mosquito borne diseases from viruses and parasites such as malaria or filariasis for analysis at the level of molecular interactions.

**Fig. 1:**
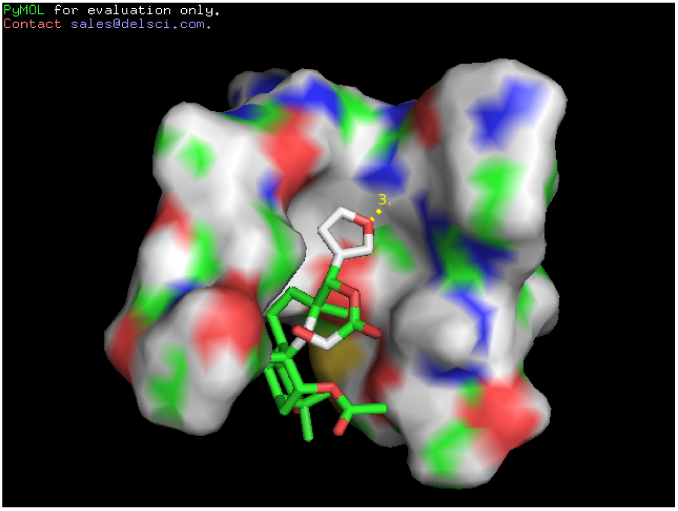
A pymol rendering of binding of the Gedunin small molecule shown in stick model to the receptor site of JAK/STAT Human Interleukin 5 protein (PDB:3VA2) shown in surface model. The dotted line indicates a hydrogen bond between oxygen atom of the ligand to a nitrogen of the ARG266 residue of a ligand binding site in the protein.

## Conclusions and future directions

We have carried out a computational experiment of analyzing the potential effect of small molecules of Neem on Dengue virus infection through simulation of molecular binding at structural level. We report here that overall contents of the Neem can be potentially useful against Dengue virus and two molecules in particular, Gedunin and Pongamol have strong interactions with important proteins of virus and its required human proteins. We report that the human proteins responsible for attachment of virus to the host cell and viral proteins themselves are blocked by some of the Neem molecules. Our research sets a ground for planning specific experimental studies for effect of Gedunin and Pongamol on control of viral infection. Present work also indicates exploring potential anti-viral applications of naturally occurring substances on mosquito borne diseases.

## ACKNOWLEDGMENTS

We would like to thank Prof. Dr. Narahari of Computer Science and Automation department of Indian Institute of Science, Bangalore for encouragement, technical review of the manuscript and providing computational facilities to carry out the research. We would also like to thank Mr. Advait Kunte for helping with software installations.

## References

1. M M Alen and D Schols. Dengue virus entry as target for antiviral therapy. J Trop Med, 2012:628475–628475, 2012.

2. F C Bernstein, T F Koetzle, G J Williams, E F Meyer, M D Brice, J R Rodgers, O Kennard, T Shimanouchi, and M Tasumi. The protein data bank: a computer-based archival file for macromolecular structures. Arch Biochem Biophys, 185(2):584–591, Jan 1978.

3. K Biswas, I Chattopadhyay, R K Banerjee, and U Bandyopadhyay. Biological activities and medicinal properties of neem (azadirachta indica). Indian Academy of Sciences, 82, 2002.

4. S B Halstead. Dengue virus-mosquito interactions. Annu Rev Entomol, 53:273–291, 2008.

5. J J Irwin and B K Shoichet. Zinc–a free database of commercially available compounds for virtual screening. J Chem Inf Model, 45(1):177–182, Jan-Feb 2005.

6. R Maheswaran and S Ignacimuthu. A novel herbal formulation against dengue vector mosquitoes aedes aegypti and aedes albopictus. Parasitol Res, 110(5):1801–1813, May 2012.

7. D Mairiang, H Zhang, A Sodja, T Murali, P Suriyaphol, P Malasit, T Limjindaporn, and R L Finley. Identification of new protein interactions between dengue fever virus and its hosts, human and mosquito. PLoS One, 8(1), 2013.

8. G M Morris, R Huey, W Lindstrom, M F Sanner, R K Belew, D S Goodsell, and A J Olson. Autodock4 and autodocktools4: Automated docking with selective receptor flexibility. J Comput Chem, 30(16):2785–2791, Dec 2009.

9. N T Ngo, X T Cao, R Kneen, B Wills, V M Nguyen, T Q Nguyen, V T Chu, T T Nguyen, J A Simpson, T Solomon, N J White, and J Farrar. Acute management of dengue shock syndrome: a randomized double-blind comparison of 4 intravenous fluid regimens in the first hour. Clin Infect Dis, 32(2):204–213, Jan 2001.

10. M M Parida, C Upadhyay, G Pandya, and A M Jana. Inhibitory potential of neem (azadirachta indica juss) leaves on dengue virus type-2 replication. J Ethnopharmacol, 79(2):273–278, Feb 2002.

11. I A Rodenhuis-Zybert, J Wilschut, and J M Smit. Dengue virus life cycle: viral and host factors modulating infectivity. Cell Mol Life Sci, 67(16):2773–2786, Aug 2010.

12. D M Watts, D S Burke, B A Harrison, R E Whitmire, and A Nisalak. Effect of temperature on the vector efficiency of aedes aegypti for dengue 2 virus. Am J Trop Med Hyg, 36(1):143–152, Jan 1987.

